# The Prognostic Value of Tumor-Infiltrating Immune Cells in Gynecologic Cancers

**DOI:** 10.1101/2021.01.31.429066

**Authors:** Waichung Chen, Tuo Hu, Chunbo He

## Abstract

Immunotherapy has changed the standard of treatment for many cancers. However, only a small number of gynecologic cancer patients benefit from immunotherapy. The intra-tumoral immune landscapes are suggested as a predictor of the response to immunotherapies, but there are no studies that provide a comprehensive immune characterization for gynecologic cancers. To characterize cellular compositions of the immune infiltrates and investigate if the immune landscape is a predictor for patient prognosis in gynecologic cancers, we analyzed tumor immune infiltrates of ovarian cancer, cervical cancer, and uterine cancer from The Cancer Genome Atlas Program (TCGA) using QuanTIseq and EPIC. Ovarian cancer had the highest percentage of total immune cells. Cervical cancer and uterine corpus endometrial carcinoma have lower percentages of immune cells with 17% and 16%, respectively. Furthermore, ovarian cancer had a significantly higher monocyte and M2-liked macrophage percentage, but a lower percentage for CD8 T cells and neutrophils compared to cervical cancer and uterine cancer. Cervical cancer had the highest percentage for M1-liked macrophages and the lowest for CD4 T cells. Uterine cancer had the highest percentage of dendritic cells. In cervical cancer, higher cell infiltration of CD8 T-Cells and M2-liked macrophages was associated with a better prognosis. In uterine cancer, patients with a higher number of dendritic cells and CD8 T-Cells had significantly better clinical outcomes. However, higher CD4 T-cell infiltration was associated with a poor prognosis in uterine cancer. Interestingly, the patient survival was not affected by the infiltration of any individual immune cells which we analyzed in ovarian cancer. We identified and validated four immune subtypes associated with distinct immune cell infiltration in gynecologic cancers. Cervical and uterine cancer patients from an immune-desert subtype that had the least amount of lymphocyte infiltration and a high level of monocyte had the worst prognosis. By contrast, cervical and uterine cancer patients from an immune-warm subtype that had higher infiltration of CD8 T-cell, natural killer (NK) cells, and dendritic cells (DCs) had the best prognosis. However, the survival rate of ovarian cancer patients is similar among the four different subtypes. Our study provides a conceptual framework to understand the tumor immune microenvironment of different gynecologic cancers.

## 1. Introduction

Gynecologic cancer arises from women’s reproductive system and is of five main types: cervical cancer, ovarian cancer, uterine cancer, vaginal cancer, and vulvar cancer. The three most common types (cervical cancer, ovarian cancer, and uterine cancer) account for 14.5% of the 8.6 million estimated new cancer cases, and 14.1% of the 4.2 million new cancer deaths among women worldwide in 2018 [1]. Except for ovarian cancer, gynecologic cancers generally have a relatively good prognosis if the cancers are detected in the early stages [2,3]. However, there are limited treatment options for patients with advanced stages or recurrent gynecologic cancers because these cancers retain a generally very poor response to current standard therapies [4]. Therefore, the development of new therapeutic approaches to treat gynecologic cancer is urgently needed.

Cancer therapy has recently experienced a dramatic revolutionary change because of the appearance of immunotherapy as a standard treatment option for several malignancies [5]. In particular, adoptive T cell therapy (ACT), e.g., with CAR-T cells, and immune checkpoint blockade (ICB), e.g., anti-PD1/PD-L1 antibodies, have demonstrated durable response and unparalleled clinical benefit in a subset of patients across multiple types of tumor [6]. Many studies have been established trying to apply immunotherapy to treat gynecologic cancer [7]. U.S. Food and Drug Administration (FDA) has approved ICB-based treatments for a subset of patients with cervical cancer and uterine cancer, but only a small number of patients, such as those with mismatch repair (MMR) deficiency, respond well to the ICB immunotherapy [8]. Clinical trials as well as preclinical studies showed that current standard immunotherapies did not have satisfactory results in most gynecologic cancer patients, indicating the existence of intrinsic immune resistance in the tumors [9]. Thus, understanding the mechanisms that support the response and resistance to the treatments will provide a way to design rational combination strategies using immunotherapy.

In developing immunotherapies and investigating resistance, the tumor immune microenvironment came into focus. Studies showed that the tumor-immune cell interaction played a critical role in regulating cancer progression. Depending on the cellular composition of the intra-tumoral immune infiltrates, immune cells can either promote or control tumor growth [10]. Using TCGA data and bioinformatics tools, recent studies characterize the immune microenvironment in pan-cancer or cancer-specific settings and their association with patient survival [11]. However, there are no studies that provide a comprehensive immune characterization specifically for gynecologic cancers.

To have a better understanding of the tumor immune microenvironment and its role in cancer progression in gynecologic cancer, in the present study we characterized cellular compositions of the immune infiltrates in cervical cancer, uterine cancer, and ovarian cancer. We identified four immune subtypes according to the immune landscape and analyzed the association of immune subtypes with patients’ prognosis in these cancers.

## 2. Results

### 2.1. The Landscape of Immune Infiltration in Gynecologic Cancers

To analyze the immune cell subtype composition in the tumors of gynecologic cancers, the upper quartile normalized RSEM (RNA-Seq by Expectation Maximization) mRNA-seq data from 1063 TCGA tumor samples (CESC: Cervical and endocervical cancers, *n* = 285; OV: Ovarian serous cystadenocarcinoma, *n* = 256; UCEC: Uterine Corpus Endometrial Carcinoma, *n* = 522) were collected from the Broad Institute FireBrowse portal (http://gdac.broadinstitute.org). By performing QuanTIseq analysis of these tumors based on the mRNA gene expression profile, the proportions of 10 types of infiltrating immune cells were characterized (Figure 1A). Interventionary studies involving animals or humans, and other studies require ethical approval must list the authority that provided approval and the corresponding ethical approval code.

**Figure 1.**
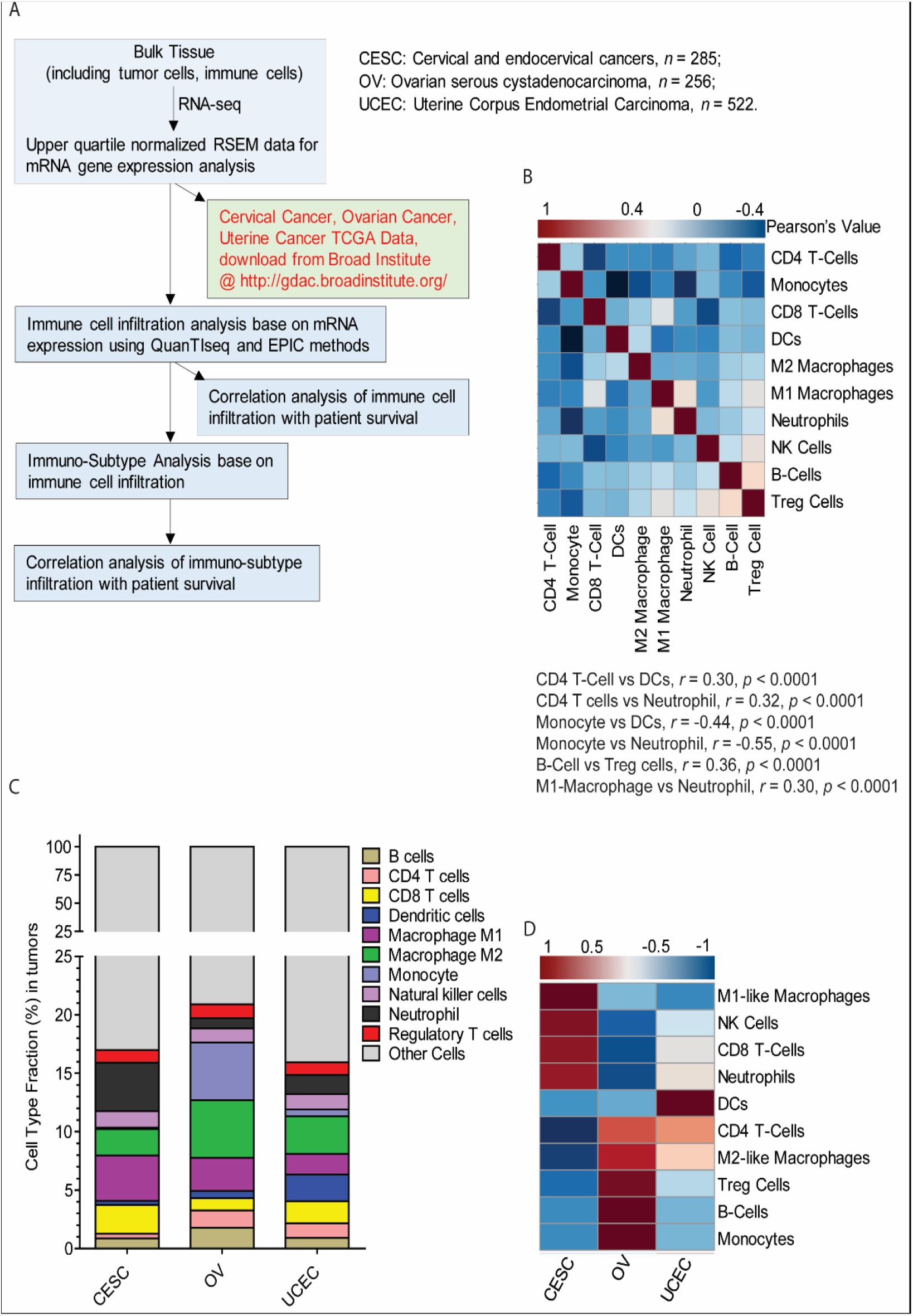
The overall landscape of immune infiltration in gynecologic cancers. (**A**) Flowchart of the conducted study process of the stages and methods used. (**B**) Correlation analysis among individual immune infiltrates in gynecologic cancers. The blue color represents a trend of negative correlation, and the red color indicates a trend of positive correlation. (**C**) Average fraction of each immune infiltrates in ovarian, cervical, and uterine cancers. (**D**) Comparison Heatmap of the average proportion of immune infiltrates in ovarian, cervical, and uterine cancers. The blue color represents a relatively lower infiltrated fraction, and the red color indicates a relatively higher infiltrated fraction. RSEM: RNA-Seq by Expectation Maximization; TCGA: The Cancer Genome Atlas Program; NK: Natural Killer Cells; DCs: dendritic cell.

The proportions of different tumor infiltrated immune cell subpopulations were weakly to moderately correlated (Figure 1B). Notably, the infiltration of CD4 T cells (non-regulatory CD4+ T cells) was positively correlated with dendritic cell (DCs) population (*r* = 0.30, *p* < 0.0001), while the fraction of Treg cells (regulatory CD4+ T cells) was positively correlated with the proportion of B cells. As shown in Figure 1C, ovarian cancer had a relatively higher average percentage of immune cells, at approximately 21%, while cervical cancer and uterine corpus endometrial carcinoma had similar percentages of total immune cells with 17% and 16%, respectively (Figure 1C). The heatmap in Figure 1D indicate the relative average proportions of these 10 immune cells. In ovarian tumors, there were relatively higher infiltrated monocytes, B cells, Treg cells, and M2-liked macrophages, but lower proportions of NK cells, CD8 T cells, and neutrophils compared to cervical cancer and uterine cancer. Cervical cancer had higher infiltrated levels for M1-liked macrophage, NK cells, and CD8 T cells, but the lowest CD4 T cells. Uterine carcinoma had a relatively higher percentage of dendritic cells compared to the other two tumors (Figure 1D).

### 2.2. The Fraction of Individual Type of Immune Cells in Gynecologic Cancers

To further characterize the features of the fraction of infiltrated individual type of immune cells in gynecologic cancers, the absolute proportions of each of the 10 immune cells were compared among ovarian, cervical, and uterine cancers. Among these three cancers, ovarian cancer had the highest proportion of total immune cells, and the highest level of M2-liked macrophages and B cells, but the lowest ratio of CD8 T cells (Figure 2A–I). Cervical cancer had the highest infiltrated cellular level for neutrophils, but the lowest level of M2-liked macrophages and CD4 T cells. Uterine cancer had a significantly higher level of infiltrated DCs than ovarian cancer and cervical cancer (Figure 2A–I).

**Figure 2.**
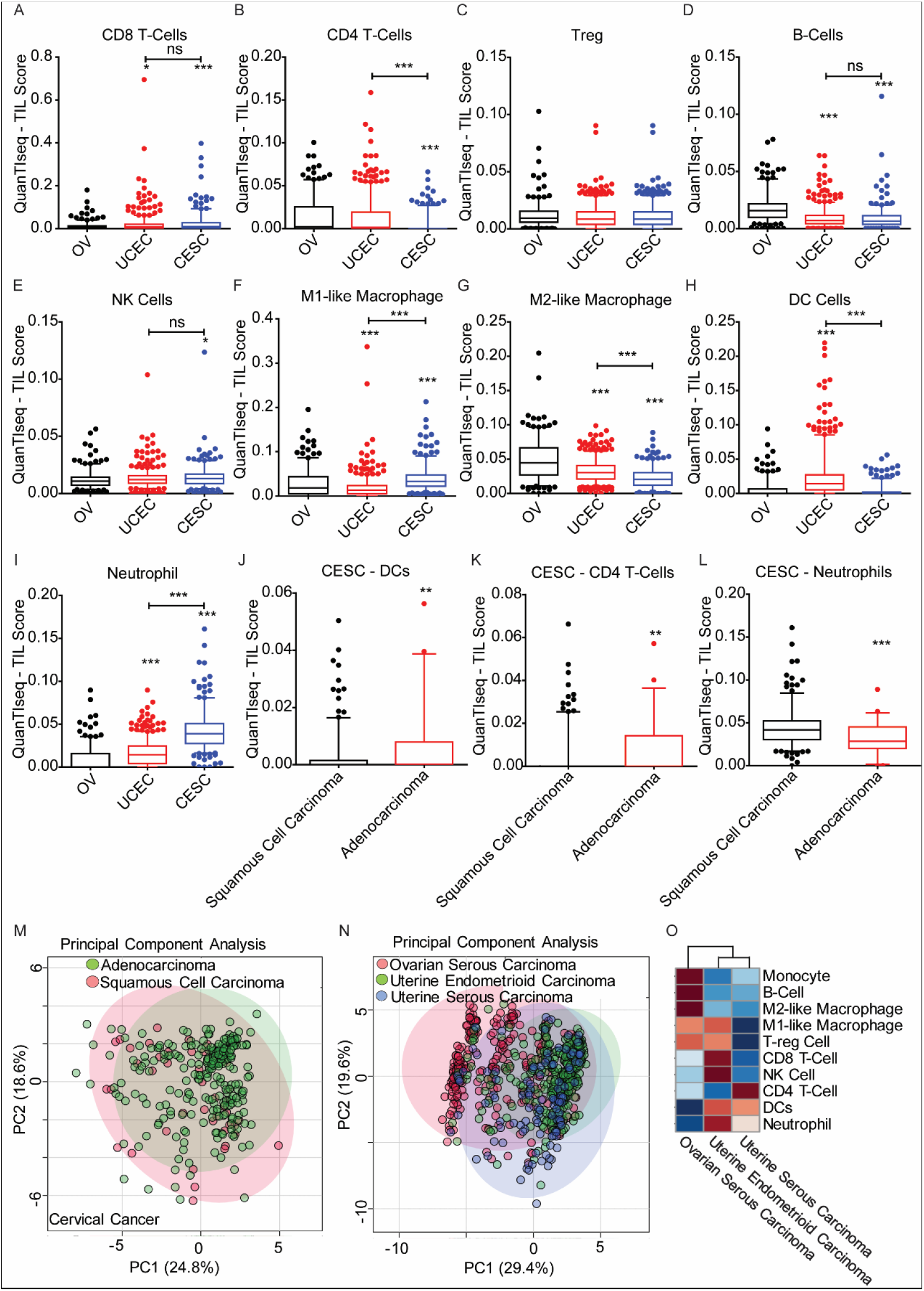
Comparison of tumor immune infiltrates among ovarian, cervical, and uterine cancers. (A– I) Comparison of significantly different percentages of tumor immune infiltrates for ovarian (*n* = 256), cervical (*n* = 285), and uterine (*n* = 522) cancers. One-way ANOVA with Tukey’s post-test was performed to analyze differences in the fraction of infiltrated immune cells among three cancers. *p* values < 0.05 were considered as statistically significant. *: *p* < 0.05; **: *p* < 0.01; ***: *p* < 0.001. **(J–L)** Comparison of tumor immune infiltrates in two subtypes of cervical cancer: cervical squamous cell carcinoma (*n* = 248) and cervical adenocarcinoma (*n* = 46). Three (DCs, non-regulatory CD4 T cells, and neutrophils) of ten immune cells demonstrated significant differences between the two subtypes. *T*-Test with unequal variances and two-tailed were performed to analyze the differences between the two subtypes. *p* values < 0.05 were considered as statistically significant. *: *p* < 0.05; **: *p* < 0.01; ***: *p* <0.001. **(M)** Scatterplot showing the distribution of cervical squamous cell carcinoma and cervical adenocarcinoma using unsupervised principal component analysis (PCA) based on the fraction of ten infiltrating immune cells. (N) Scatterplot showing the distribution of ovarian serous carcinoma (*n* = 256), uterine endometrioid carcinoma (*n* = 399), and uterine serous carcinoma (*n* = 109) using unsupervised principal component analysis (PCA) based on the fraction of ten infiltrating immune cells. **(O)** Heatmap with consensus clustering to compare the composition and proportion of tumor immune infiltrates in ovarian serous carcinoma, uterine endometrioid carcinoma, and uterine serous carcinoma. The blue color represents a relatively lower infiltrated fraction, and the red color indicates a relatively higher infiltrated fraction. CESC: Cervical and endocervical cancers, *n* = 285; OV: Ovarian serous cystadenocarcinoma, *n* = 256; UCEC: Uterine Corpus Endometrial Carcinoma, *n* = 522; NK: Natural Killer Cells; DCs: Dendritic Cells.

There are two different pathologic types for cervical cancer, cervical adenocarcinoma and cervical squamous cell carcinoma [12]. Cervical adenocarcinoma had significantly higher infiltrated DCs and CD4-T cells, but lower neutrophils than cervical squamous cell carcinoma (Figure 2J–L). A scatterplot of the principal component analysis (PCA) based on the fraction of 10 immune cells showed that the sample plots of squamous cell carcinoma and adenocarcinoma did not cluster to different groups, but crossed together (Figure 2M), indicating that the immune landscapes of cervical adenocarcinoma and cervical squamous cell carcinoma were similar overall.

Uterine serous carcinoma shares many pathologic features with ovarian serous carcinoma [13]. To investigate if they also have similar immune cell composition in their tumors, unsupervised principal component analysis (PCA) was performed on ovarian serous carcinoma, uterine serous carcinoma, and uterine endometrioid carcinoma. As shown in Figure 2N, uterine serous carcinoma and uterine endometrioid carcinoma plots clustered together relatively and separated from ovarian serous carcinoma, indicating that the immune landscape of uterine serous carcinoma is more like that of uterine endometrioid carcinoma rather than of ovarian serous carcinoma. Consensus clustering-based analysis also confirmed the similarity of the immune cell infiltration landscape in these two subtypes of uterine cancer (Figure 2O).

### 2.3. Clinical Characteristics and Prognostic Value of the Infiltrated Immune Cells

Recent studies have reported a significant correlation between the tumor infiltrated immune cells and their prognostic role in different types of cancers [14]. For example, a high peritumoral density of CD8 T cells, B cells, and mature DCs were associated with longer survival, while higher infiltration of neutrophil granulocytes predicted a poor prognosis in melanoma [15].

To evaluate the prognostic value of individual infiltrating immune cells in gynecologic cancers, we analyzed the patients’ overall and progression-free survival rate, relating this to the fraction of infiltrated immune cells in ovarian, cervical, and uterine cancers. Interestingly, although ovarian cancer had the highest density of immune cells among these three cancers, none of the immune cell subtypes were significantly associated with patient survival (Figure 3A and Figure A1).

**Figure 3.**
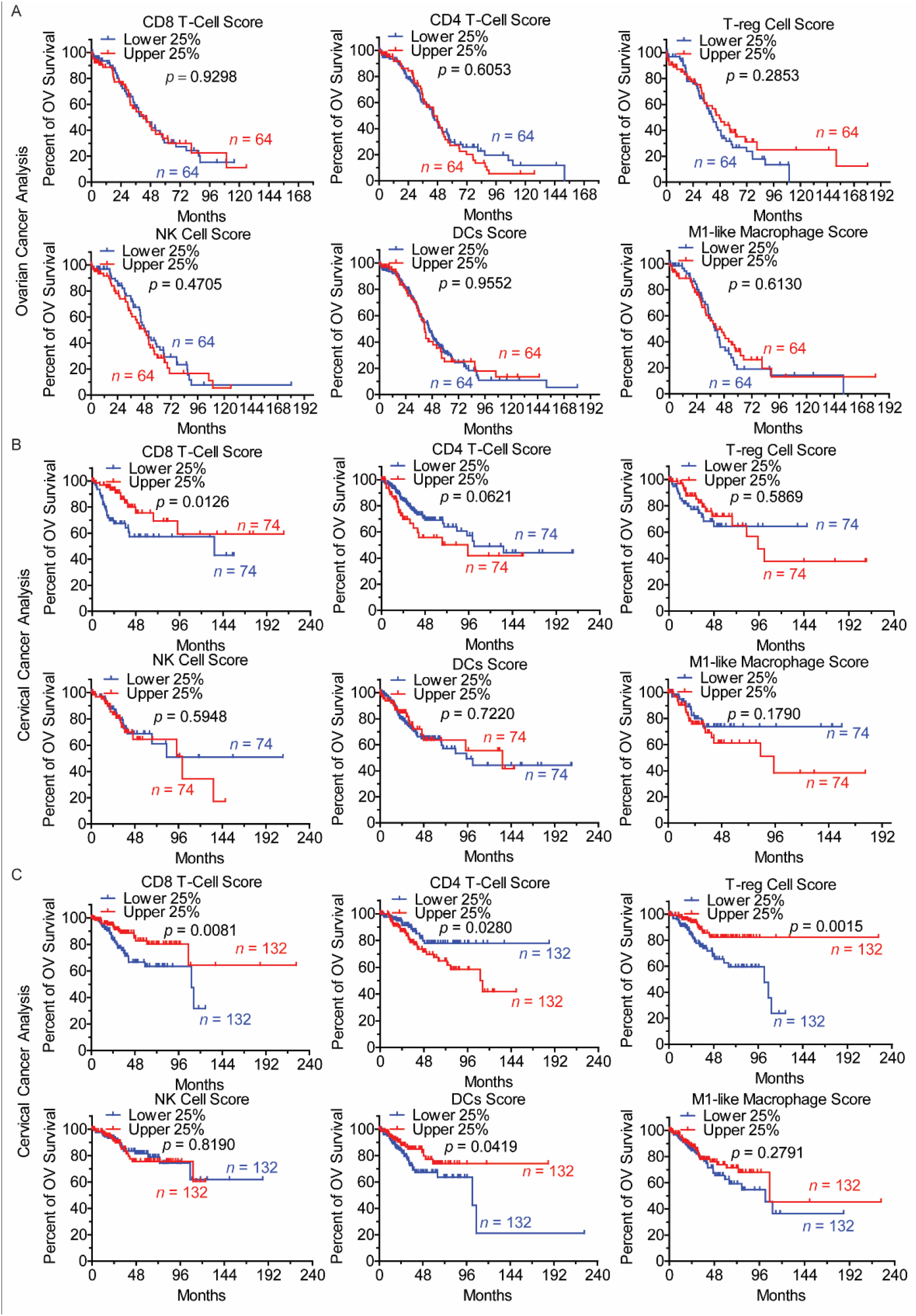
Prognostic value of tumor-infiltrating immune cells in gynecologic cancers. (A) Kaplan– Meier curves showing the overall survival (OS) and progression-free survival (PFS, in Supplementary Figure A1) of ovarian cancer patients stratified by the proportion of infiltrated immune cells. **(B)** Kaplan–Meier curves showing the overall survival (OS) and progression-free survival (PFS, in Supplementary Figure A2) of cervical cancer patients stratified by the proportion of infiltrated immune cells. **(C)** Kaplan–Meier curves showing the overall survival (OS) and progression-free survival (PFS, in Supplementary Figure A3) of uterine cancer patients stratified by the proportion of infiltrated immune cells. For each specific immune cell, survival data of samples with higher (upper 25%) and lower (lower 25%) immune cell fraction were compared using Kaplan-Meier survival estimates. *P*-value was calculated by the log-rank test. *p* values < 0.05 were considered as statistically significant. OV survival: overall survival; NK: natural killer cells; DCs: dendritic cells.

Cervical cancer had a relatively higher average level of infiltrated CD8 T cells than ovarian and uterine cancer (Figure 1C, D). Since CD8+ T cells can directly kill tumor cells via cytotoxic molecules, such as granzymes and perforin, it is the major target cell for cancer immunotherapy. We found that a higher abundance of CD8 T cells was associated with a better prognosis in cervical cancer (Figure 3B), suggesting that CD8 T cells play a critical role in controlling cervical cancer progression. In the tumor microenvironment, M2-like macrophages are usually considered as one of the compositions which can promote tumor growth, but cervical cancer patients with a higher fraction of M2-like macrophage in their tumor had a significantly better overall and progression-free survival rate (Figure A2). For the remaining other immune cells, their infiltration did not significantly affect patients’ survival in cervical cancer (Figures 3B and A2).

Compared to ovarian and cervical cancer, the prognosis of uterine cancer was associated with different immune cell types. As in cervical cancer, a higher CD8 T cell infiltration also predicted a better overall and progression-free survival rate in uterine cancer (Figures 3C and A3). A higher density of DCs was also correlated with longer overall and progression-free survival (Figures 3C and A3). As an immunosuppressive subset of CD4+ T cells, regulatory T (Treg) cells were reported as suppressing antitumor immunity, thereby hindering immunosurveillance against cancer development in many types of cancer. However, we found that the fraction of infiltrated Treg cells was positively associated with the rates of overall and progression-free survival in uterine cancer (Figures 3C and A3). In contrast, the infiltration of non-regulatory CD4+ T cells was negatively correlated with patient prognosis (Figures 3C and A3), indicating that Treg cells may play a dual role in regulating uterine cancer progression.

### 2.4. The Fraction of Individual Types of Immune Cells in Gynecologic Cancers

Due to the heterogeneity of immune cell infiltration in tumors, we tried to cluster the samples based on the fraction of infiltrated immune cell subtypes. Unsupervised PCA analysis showed that the tumor sample scatterplots were predominantly separated into four clusters: immuno-subtype I to immuno-subtype IV (Im-munoSub-I, II, III, and IV) (Figure 4A,B). As shown in Figure 4C, there were only a few cervical cancer samples grouped into ImmunoSub-I with ovarian cancer being in the majority, counting for 70%. However, ovarian cancer accounted for less than 10% in ImmunoSub-III and ImmunoSub-IV. Uterine cancer counted for around 65% in ImmunoSub-IV (Figure 4C).

**Figure 4.**
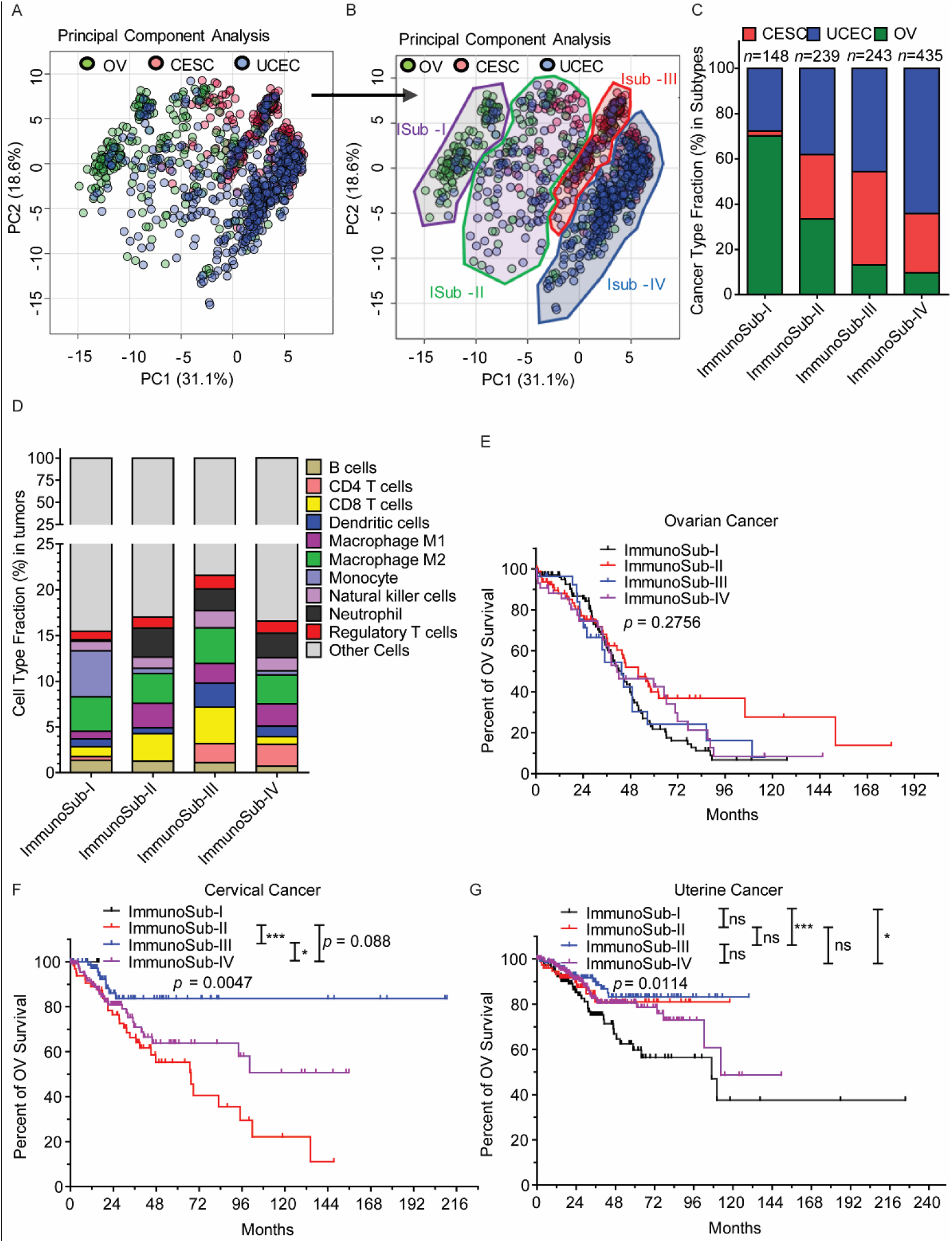
The overall landscape of immune infiltration in gynecologic cancers. (A) Scatterplot showing the distribution of all patients with gynecologic cancers using unsupervised principal component analysis (PCA) based on the fraction of ten infiltrating immune cells. (B) Immune subtypes were arranged based on the distribution of patients’ scatterplot. Each plot represented a patient. Patients were clustered into four immuno-subtypes, ImmunoSub-I, II, III, and IV (ISub-I, II, III, IV). (C) The distribution of immune subtypes among ovarian, cervical, and uterine cancers. (D) Tumor immune infiltration patterns of ten immune cells across four immune subtypes. The column height of each immune cell means the average faction of infiltrated cells. (E–G) Kaplan–Meier curves showing the overall survival (OS) of all ovarian cancer (E), cervical cancer patients (F), and uterine cancer (G) stratified by the immune subtypes. Cervical cancer patients stratified by the proportion of infiltrated immune cells. *P*-value was calculated by the log-rank test among subtypes. *p* values < 0.05 were considered as statistically significant. ns: no significant difference; *: *p* < 0.05; **: *p* < 0.01; ***: *p* < 0.001. OV survival: overall survival; OV: ovarian cancer; UCEC: Uterine Corpus Endometrial Carcinoma; CESC: Cervical and endocervical cancers.

Regarding the fraction of different cell types in the four subtypes, the average fraction of M2-like macrophage remains similar in all four groups (Figure 4D). ImmunoSub-I could be considered as the immune cold group because it had the least amount of total immune cells, but it has the greatest percentage of monocyte. Tumors in ImmunoSub-II had the lowest infiltrated non-regulatory CD4+ T cells (Figure 4D). Although ImmunoSub-III tumors contain the highest percentage of total immune cells, there were fewer monocytes. Tumors from ImmunoSub-III had a relatively higher fraction of CD8 T cells, DCs, CD4 T cells, and NK cells (Figure 4D). Subtype IV has the highest proportion of CD4 T cells and the least fraction of CD8 T cells (Figure 4D).

Kaplan-Meier survival estimates with a log-rank test were used to compare the survival difference among four immune subtypes. As shown in Figure 4E–G, the survival difference of the subtypes was dependent on cancer type. We did not observe a significant prognostic impact of the immune subtypes in ovarian cancer (Figure 4E). However, unlike ovarian cancer, there was a significant difference in the survival rate among the three subtypes, ImmunoSub-II, III, and IV. (There were only three cervical cancer patients in ImmunoSub-I) (Figure 4F). Overall, cervical cancer patients from ImmunoSub-III Subtype had the best survival rate for both overall survival and progression-free survival. Although worse than ImmunoSub-III, ImmunoSub-IV was associated with a better prognosis compared to ImmunoSub-II (Figure 4F). For uterine cancer, the Kaplan-Meier survival curve showed that the four subtypes also had a significant difference in survival rate (Figure 4G). As in cervical cancer, the immune-warm group ImmunoSub-III was associated with the best prognosis for both overall survival and progression-free survival. By contrast, the immune-cold subtype ImmunoSub-II was associated with the worst outcomes among all subtypes in uterine cancer. The remaining ImmunoSub-I and IV had an intermediate prognosis (Figure 4G).

## 3. Discussion

Oncology has recently experienced a dramatic revolutionary change because of the appearance of immunotherapy as an option for cancer treatment. Immune checkpoint inhibitors (ICIs) and chimeric antigen receptor (CAR) T-Cell Therapy have been transformative in both solid tumors and hematologic malignancies [16]. Patients with previously terminal illnesses, including relapsed or refractory B-cell lymphoma, advanced non-small-cell lung cancer (NSCLC) and metastatic melanoma, have experienced profound responses and unprecedented clinical benefits [17–19]. Unfortunately, response and survival benefits were only limited to a small subset of patients with gynecologic cancers [9]. So far, none of the single-agent-based checkpoint immunotherapies or adoptive engineering immune cell therapies have demonstrated a statistically significant and clinically meaningful improvement in overall survival as compared with current standard treatments in ovarian cancer [8]. Monotherapy of ICIs only showed a moderate overall response rate in mismatch repair (MMR) deficiency and microsatellite instability (MSI) high endometrial cancer [20], which may count for around 30% of all endometrial tumors [21]. Although pembrolizumab (targets PD-1) was approved by the FDA for patients with recurrent or metastatic cervical cancer with disease progression on or after chemotherapy whose tumors have a high level of PD-L1, response to pembrolizumab alone has shown only modest clinical benefits [22]. Results of these clinical trials and preclinical investigations suggest that a potently immunosuppressive microenvironment exists in gynecological tumors. Thereby a better understanding of interactions between the tumor cells, immune cells, and other components of the tumor microenvironment may provide ways to improve the prediction of who could benefit from immunotherapies and help the development of the next generation of immunotherapies.

In this study, we presented a comprehensive characterization of tumoral-infiltrated immune cell subtypes in ovarian, cervical, and uterine cancers. We also identified four immune subtypes based on the proportions and compositions of infiltrated immune cells. We found that each of the gynecologic cancers showed distinct immune infiltrate profiles, and the prognostic value of tumor-infiltrating immune cells and immunosubtypes depended on cancer types. Our study provides a conceptual framework to understand the relationships between therapeutic response and immune infiltrates in gynecologic cancers and may also demonstrate clinical implications for personalized immunotherapy.

Generally speaking, the tumor immune infiltrate can be classified as a ‘pro-inflammatory’ phenotype, in which the tumors have higher infiltrating T cells, DCs, M1-like macrophages, and with an immune active cytokine profile; and an ‘immunosuppressive’ phenotype, in which the tumors have more inhibitory immune cells, such as Treg cells, M2-like macrophages, and myeloid-derived suppressor cells (MDSCs) [23]. A large number of studies have supported that immunosurveillance is a significant contributor in preventing the development and progression of malignancies [24]; therefore, in a ‘pro-inflammatory’ tumor microenvironment, for example, increased infiltrating CD8 T lymphocytes are associated with improved clinical outcomes in most cases [25]. We found that higher CD8 T infiltrate indeed predicted a longer survival in both cervical and uterine cancers (Figure 3B,C). Besides, cervical and uterine cancer patients from the ImmunoSub-III, an immune-warm subtype that demonstrated the highest levels of infiltration by pro-inflammatory effectors including CD8 T cells, DCs, and NK cells, had the best prognosis when compared to patients from other subtypes (Figure 4F,G). These results indicating the cytotoxicity of CD8 T cells may play a critical role in controlling disease progression during treatment, whether immunotherapy or other therapies in these cancers. These results also suggest that a combination of standard treatments with immune activation reagents, for example, cytokines for inducing CD8 T/NK cell enrichment, may lead to improved clinical outcomes.

Ovarian cancer is generally considered as “cold tumor”[26]. However, according to our results (silico-based analysis), ovarian cancer had the lowest CD8+ T cells and NK cells among these three cancers, but had the highest level of B cells and monocytes. Ovarian cancer also had the highest fraction of total immune cells (Figures 1 and 2). However, in contrast to cervical cancer and uterine cancer, infiltrating CD8 T cells, as well as other immune cells, failed to show a significant effect on overall and progression-free survival in ovarian cancer (Figures 3A and A1). Ovarian cancer patients from different classified immune subtypes also showed similar survival curves (Figure 4E). These results suggest that immune cells might be dysregulated in ovarian cancer. Our results were also supported by increasing evidence which indicates that ovarian cancer builds a potently immunosuppressive microenvironment via interaction of complex networks of regulatory factors, and single ICB agent-based immunotherapy cannot control tumor growth in ovarian cancer. Therefore, more studies need to be conducted to draw an integrated immunosuppressive network map in ovarian cancer. On top of this, multitarget-based therapeutic strategies that modulate multiple immunosuppressive signals should lead to improved clinical outcomes using current therapies.

Unlike the ‘pro-inflammatory’ phenotype, tumors with an ‘immunosuppressive’ phenotype contain more inhibitory immune cells, such as M2-like macrophages and Treg cells [27]. Since a major role of these cells is to disrupt the normal functions of other effector cells, the ‘immunosuppressive’ phenotype is usually considered as an unfavorable factor in prognosis [28–30]. Perhaps counterintuitively, we found that infiltrating M2-like macrophages were associated with a better survival rate in cervical cancer (Figure A2), and a higher fraction of infiltrated Treg predicted a longer survival time in uterine cancer patients (Figure 3C). In fact, this phenomenon is not limited to cervical and uterine cancer, and favorable outcomes have also been observed in higher tumoral infiltration by Treg cells in pancreatic, colorectal, bladder, and esophageal cancers [31–33]. These results indicate a complex cellular cross-talking within the tumor microenvironment. For example, Zhang et al. reported that Treg depletion failed to relieve immunosuppression; instead, it led to accelerated tumor progression because of reprograming of the fibroblast population in pancreatic cancer [33]. Another notable point is that there is always a feedback loop between immune activation signals and immunosuppressive signals [27], and an existing anti-inflammatory response may indicate increased tumoral immunogenicity. Therefore, the role of immunosuppressive subtype cells during the onset and progression of gynecological cancers, as well as in the response of cancer treatments, need to be further investigated.

In recent TCGA studies specifically focused on gynecological cancers, researchers classified tumor samples into different subtypes based on their genomic features, transcriptional levels, or epigenetic modifications. However, these analyses showed limited information about the immune landscape of these cancers [34– 36]. Our analysis was also based on transcriptional data of genes reflecting various immunological processes, but we further extended the results to cellular composition and function level. These results offered a new insight into the complex immune landscape of gynecologic cancers. Our study also demonstrated that the immune infiltrate has a variable effect on the patient’s prognosis depending on the cancer type. This study also offers potential therapeutic implications for the rational design of strategies for combination therapy. For example, for ovarian cancer patients, checkpoint immunotherapies must be insufficient due to the presence of strong immunosuppressive mechanisms. This may provide clinical improvement by combining current standard-of-care therapy and using multiagent therapeutic strategies to target complex network signals and relieve immunosuppressive press. For cervical and uterine cancers in immune-warm ImmunoSub-III, checkpoint inhibitor-based immunotherapies may be used to enhance the pre-existing antitumor capability of these immune cells and further improve the survival of these patients. For those cervical and uterine cancer patients with an immune-cold tumor (ImmunoSub-I), checkpoint immunotherapies could be optimally combined with immune co-stimulatory regents, such as IL15 and IL21, which may induce the enrichment of pro-inflammatory cells and trigger an antitumor immune response.

The major limitation of our study is the lack of bench work-based data support. We also noted that there were several studies that reported results contrary to our conclusion, especially in ovarian cancer [37–39]. Sato et al. and other groups reported that CD8+ tumor-infiltrating lymphocytes were associated with a favorable prognosis in ovarian cancer [37,39]. Tsuji et al. found a favorable survival is also related to the presence of tumor-reactive T cell (CD8+) clonotypes in the tumor in ovarian cancer [40]. These studies counted CD8+ T and other immune cell levels based on immunohistochemical staining of tumor tissue microarrays. However, our results came from a silico data analysis that was based on mRNA sequencing of bulk tumor tissues. The immunohistochemistry-based method might not reflect the distribution of immune cells in entire tumor tissues, while our silico analysis lacked direct evidence support. In the future, other methods such as FACS-based immune profiling with direct patients’ samples would make the conclusions more convincing. Besides, subtype classifications in this study were based on immune cell infiltration scores, and this clustering method might be not a good model to reflect the immune landscape of ovarian cancer. There were several other models, such as consensus clustering based on cancer immune gene expression signature scores [23,41], which can be further investigated in gynecologic cancers, particularly in ovarian cancer.

In summary, we analyzed ten infiltrated immune cells and provided a comprehensive landscape of the immune microenvironment in ovarian, cervical, and uterine cancers. We determined the prognostic value of a specific subtype of tumor-infiltrating immune cells for clinical outcomes in gynecologic cancers. Our results demonstrated that the effect of the immune infiltrate on the patient’s prognosis was dependent on the cancer types. We also identified four immune subtypes of gynecologic cancers with distinct immune infiltrate characteristics and clinical outcomes. Our study provides a systematic insight into the tumor immune microenvironment of different gynecologic cancers and offers a conceptual framework for the future design of rational combination treatment strategies and optimal selection of patients to improve immunotherapy outcomes.

## 4. Materials and Methods

### 3.1. Patient Information and Datasets

In this study, the mRNA gene expression data (upper quartile normalized RSEM data for batch-corrected mRNA gene expression) [42] were obtained from the Broad Institute FireBrowse portal (http://gdac.broadinstitute.org). The clinical information was obtained from the Memorial Sloan Kettering Cancer Center cBioPortal (http://www.cbioportal.org) [43]. Based on the TCGA user guideline, all data and information that we used in the present study are “Genomic Data Commons (GDC) Open Access Data.” The GDC itself places no restrictions (other than attempts at reidentification) on analysis or publication of open access data provided through the GDC Data Portal (https://gdc.cancer.gov/about-data/data-analysis-policies). That all data were from existing and de-identified public datasets, the TCGA Publication Committee ensured that our data’s status is categorized as “No restrictions; all data available without limitations.” Informed patient consent was waived for all data we used. There were 294 cervical and endocervical cancer, 255 ovarian serous cystadenocarcinoma, and 525 uterine corpora endometrial carcinoma were used for analysis (Figure 1A, Table 1, and supplementary Table S1). The raw data, processed data, and clinical data can be found in the legacy archive of the National Cancer Institute’s Genomic Data Commons (GDC) (https://portal.gdc.cancer.gov/legacy-archive/search/f).

**Table 1.** Clinical Features in Patients.

| Variables | Cervical Cancer | Ovarian Cancer | Uterine Cancer |
| --- | --- | --- | --- |
| **Numbers** | 294 | 255 | 525 |
| **Age** |  |  |  |
| < 60 years | 231 (78.6) | 144 (56.5) | 175 (55.2) |
| ≥ 60 years | 63 (21.4) | 111 (43.5) | 350 (44.8) |
| **Race** |  |  |  |
| White | 201 (68.4) | 203 (79.6) | 354 (67.4) |
| other | 57 (19.4) | 25 (9.8) | 139 (26.5) |
| NA | 38 (12.2) | 27 (10.6) | 32 (6.1) |
| **TNM stage** |  |  |  |
| I | 156 (53.1) | 0 (0) | 326 (62.1) |
| II | 69 (23.5) | 19 (7.5) | 50 (9.5) |
| III | 45 (15.3) | 204 (80) | 120 (22.9) |
| IV | 18 (6.1) | 32 (12.5) | 29 (5.5) |
| **Histological grade** |  |  |  |
| G1 | 17 (5.8) | 1 (0.4) | 97 (18.5) |
| G2 | 130 (44.2) | 27 (10.6) | 118 (22.5) |
| G3 | 115 (39.1) | 201 (78.8) | 310 (61) |
| Gx | 32 (10.9) | 26 (10.2) | 0 (0) |
| **Radiation therapy** |  |  |  |
| YES | 174 (59.2) | 2 (0.8) | 218 (41.5) |
| NO | 94 (25.5) | 225 (88.2) | 283 (53.9) |
| NA | 4 (15.3) | 28 (11.0) | 24 (4.6) |
| **Sample Location** |  |  |  |
| Primary | 294 (100) | 255 (100) | 525 (100) |

All data are shown as numbers and percentages—numbers (percentage%). Gx: No information for the histological grade. NA: No information. TNM: The TNM Classification of Malignant Tumors.

### 4.2. Analysis of Tumor-Infiltrating Immune Cells

The composition and density of immune cells in the tumor microenvironment (TME) were analyzed by QuanTIseq, a deconvolution-based method for quantifying the fractions of immune cell types from bulk RNA-sequencing data in tumor and blood samples [44]. This deconvolution-based method formulates a system of equations that describe the gene expression of a sample as the weighted sum of the expression profiles of the admixed cell types. By using least square regression, cell type fractions can be inferred given a signature matrix and the bulk gene expression. In practice, the QuanTIseq allows us to directly analyze raw RNA-seq data and to generate absolute scores that can be interpreted as cell fractions and compared both inter- and intra-sample. The QuanTIseq analysis can give the fractions of ten immune cells relative to all cells in the samples, including B cells; CD4 T-cells (non-regulatory CD4+ T cells); CD8+ T cells; DCs (dendritic cells); M1 macrophages (classically activated macrophage; M2 macrophages (alternatively activated macrophages); monocytes; neutrophils; NK cells (natural killer cells); and Treg cells (regulatory T cells). The relative fractions of six immune cells (B cells, CD4 T-cells, CD8+ T cells, monocytes, neutrophils, and NK cells) were also evaluated by EPIC, another wildly used deconvolution-based method for analyzing immune cell types from bulk tumor gene expression data [45]. We found EPIC results showed a similar trend as QuanTIseq results in all three cancers. Furthermore, survival analysis based on QuanTIseq immune cell fractions was also similar to EPIC fractions. In all figures of the present study, we presented QuanTIseq analysis results.

### 4.2. Analysis of Tumor-Infiltrating Immune Cells

The composition and density of immune cells in the tumor microenvironment (TME) were analyzed by QuanTIseq, a deconvolution-based method for quantifying the fractions of immune cell types from bulk RNA-sequencing data in tumor and blood samples [44]. This deconvolution-based method formulates a system of equations that describe the gene expression of a sample as the weighted sum of the expression profiles of the admixed cell types. By using least square regression, cell type fractions can be inferred given a signature matrix and the bulk gene expression. In practice, the QuanTIseq allows us to directly analyze raw RNA-seq data and to generate absolute scores that can be interpreted as cell fractions and compared both inter- and intra-sample. The QuanTIseq analysis can give the fractions of ten immune cells relative to all cells in the samples, including B cells; CD4 T-cells (non-regulatory CD4+ T cells); CD8+ T cells; DCs (dendritic cells); M1 macrophages (classically activated macrophage; M2 macrophages (alternatively activated macrophages); monocytes; neutrophils; NK cells (natural killer cells); and Treg cells (regulatory T cells). The relative fractions of six immune cells (B cells, CD4 T-cells, CD8+ T cells, monocytes, neutrophils, and NK cells) were also evaluated by EPIC, another wildly used deconvolution-based method for analyzing immune cell types from bulk tumor gene expression data [45]. We found EPIC results showed a similar trend as QuanTIseq results in all three cancers. Furthermore, survival analysis based on QuanTIseq immune cell fractions was also similar to EPIC fractions. In all figures of the present study, we presented QuanTIseq analysis results.

### 4.3. Identification and Visualization of Immuno-subtypes of the Tumor Samples

An unsupervised principal component analysis (PCA) [46] was performed on all analyzed samples based on the fraction of ten infiltrating immune cells as determined by quanTIseq. The analysis was based on the methods reported by Vichi et al. and Granato et al. [47,48]. Principal Component Analysis (PCA) was visualized through the built-in R functions prcomp (). Disjoint Principal Component Analysis (CDPCA) for clustering was carried out, computing the first two principal components and classifying subtypes based on the first two objects’ scores. Matrix U (Slope value for PC1 and PC2) was used to divide subgroups. The group dividing is based on the distribution of individual samples. We subjectively separated the samples into four subtypes based on their distribution. The principal components were calculated using the built-in R functions prcomp (), and the visualized first two (PC1 and PC2) projection scatterplot was built using the ggplot2 package of R function. Samples were classified into different groups based on the distribution and clustering of their plots. A total of four groups were settled from PCA results of ten infiltrating immune cells. The clusters of different subtypes were also evaluated by Monti consensus clustering method.

### 4.4. Correlation of Immune Cell Subtypes with Overall and Progression-Free Survival

Survival analysis of ovarian, uterine, and cervical cancer patients stratified by the infiltrated immune cell subtypes was based on QuanTIseq analysis results. For each specific immune cell, survival data of samples with higher (top 25%) and lower (bottom 25%) immune cell fraction were compared using Kaplan-Meier survival estimates. The Kaplan-Meier survival curve with 95% confidence intervals was computed using Prism 5 software (San Diego, CA, USA). *p*-values were calculated using the Log-Rank test comparing the survival curves of the lower and upper 25% in each immune cell group. All samples were included for the survival analysis of four immuno-subtypes. Log-Rank test was used to determine the differences between any two different subtypes.

### 4.5. Statistical Analysis

*T*-Test with unequal variances and two-tailed were performed to analyze the differences between two groups. One-way ANOVA with Tukey’s post-test was performed to analyze the differences among three or more groups. All analyses and grafts were conducted using Prism version 5. *p* values < 0.05 were considered as statistically significant.

## Author Contributions

W.C. contributed to data collection, data analysis, and manuscript preparation. T.H. contributed to the study design, data analysis, and manuscript preparation. C.H. supervised these studies and contributed to the study design, data analysis, and manuscript preparation. All authors have read and agreed to the published version of the manuscript.

## Institutional Review Board Statement

Ethical review and approval were waived for this study, due to all data were from existing and de-identified public datasets. The TCGA Publication Committee ensured that our data’s status is categorized as “No restrictions; all data available without limitations.”

## Informed Consent Statement

Informed patient consent was waived for all data we used due to all data were from existing and de-identified public datasets. The TCGA Publication Committee ensured that our data’s status is categorized as “No restrictions; all data available without limitations.”

## Data Availability Statement

The data presented in this study are openly available in TCGA database. The raw data, processed data, and clinical data can be found in the legacy archive of the National Cancer Institute’s Genomic Data Commons (GDC) (https://portal.gdc.cancer.gov/legacy-archive/search/f).

## Appendix A

**Figure A1.**
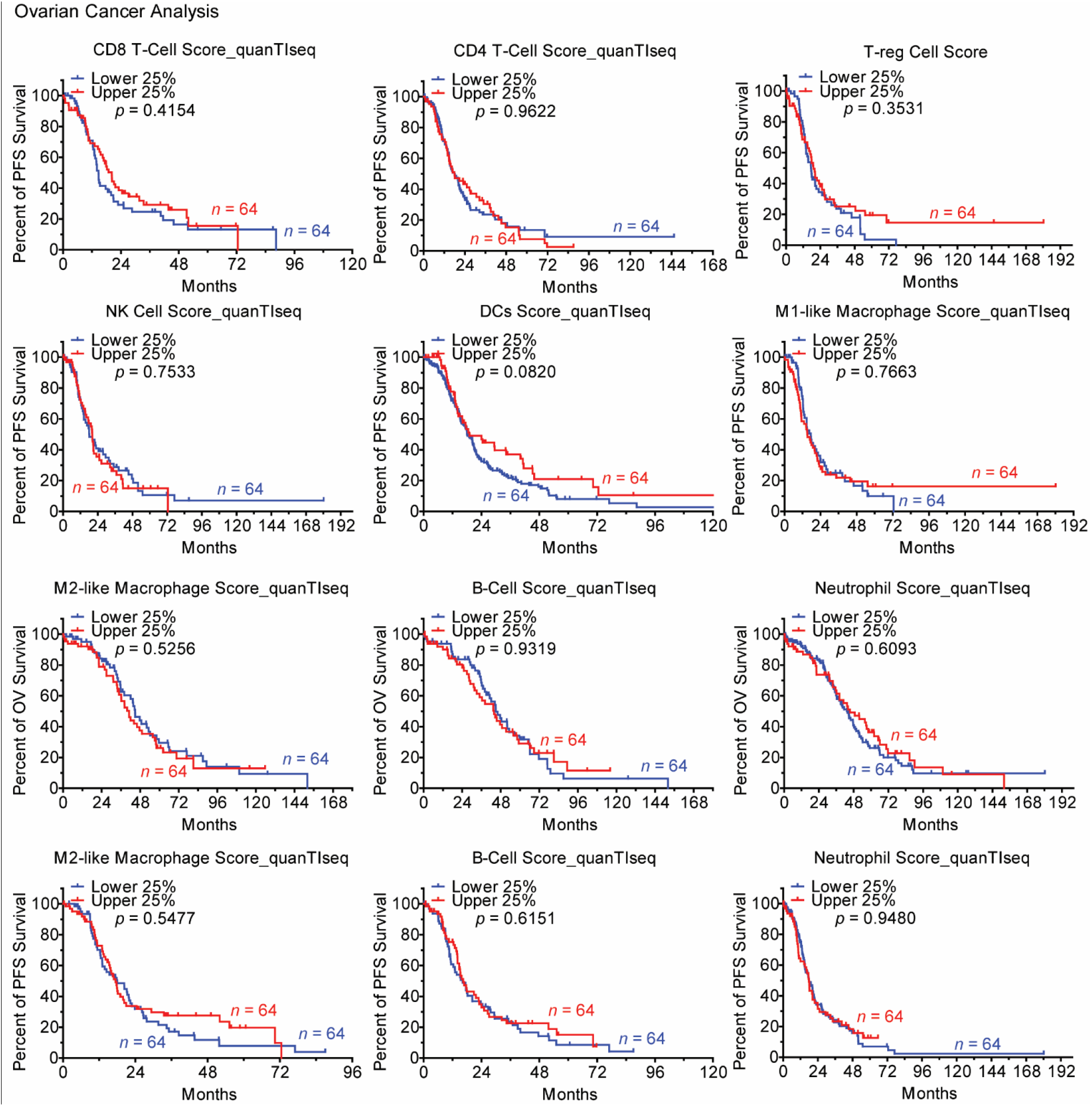
Prognostic value of tumor-infiltrating immune cells in ovarian cancers. Kaplan–Meier curves showing the overall survival (OS) and progression-free survival (PFS of ovarian cancer patients stratified by the proportion of infiltrated immune cells. For each specific immune cell, survival data of samples with higher (upper 25%) and lower (lower 25%) immune cell fraction were compared using Kaplan-Meier survival estimates. P value was calculated by the log-rank test. *p* values < 0.05 were considered as statistically significant. NK: natural killer cells; DCs: dendritic cells.

**Figure A2.**
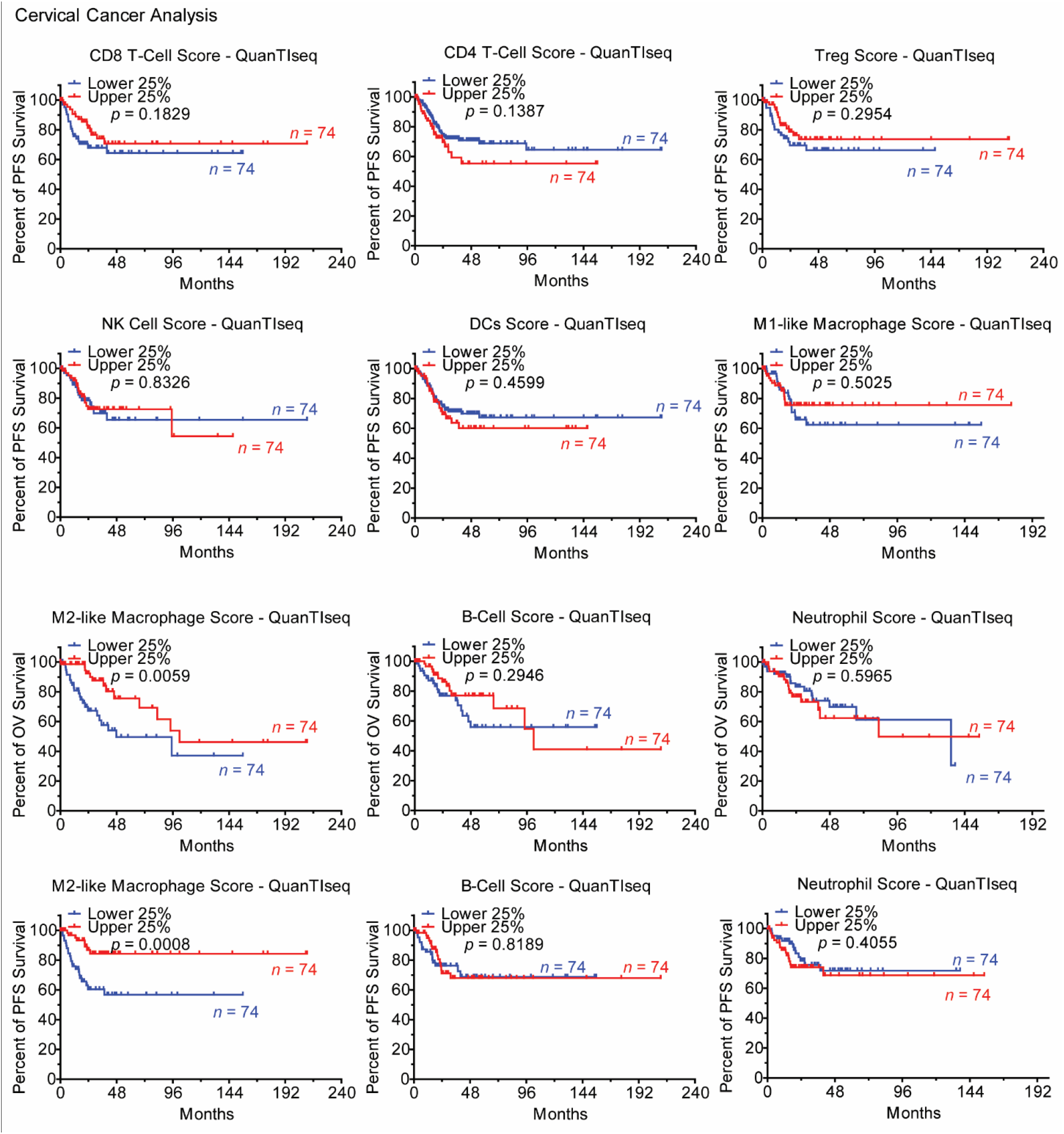
Prognostic value of tumor-infiltrating immune cells in cervical cancers. Kaplan–Meier curves showing the overall survival (OS) and progression-free survival (PFS) of cervical cancer patients stratified by the proportion of infiltrated immune cells. For each specific immune cell, survival data of samples with higher (upper 25%) and lower (lower 25%) immune cell fraction were compared using Kaplan-Meier survival estimates. P value was calculated by the log-rank test. *p* values < 0.05 were considered as statistically significant. NK: natural killer cells; DCs: dendritic cells.

**Figure A3.**
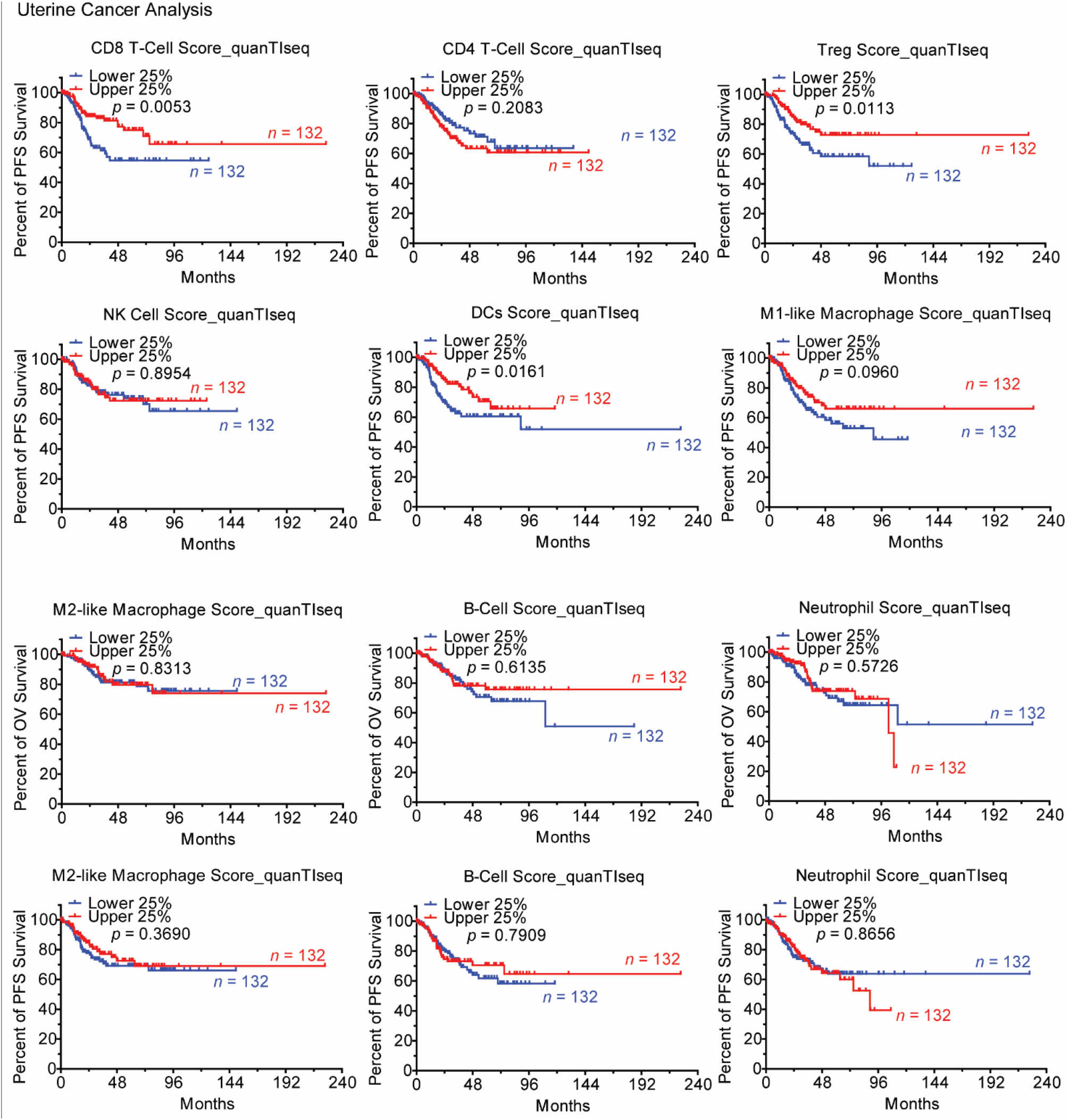
Prognostic value of tumor-infiltrating immune cells in gynecologic cancers. Kaplan–Meier curves showing the overall survival (OS) and progression-free survival (PFS) of uterine cancer patients stratified by the proportion of infiltrated immune cells. For each specific immune cell, survival data of samples with higher (upper 25%) and lower (lower 25%) immune cell fractions were compared using Kaplan-Meier survival estimates. P value was calculated by the log-rank test. *p* values < 0.05 were considered as statistically significant. NK: natural killer cells; DCs: dendritic cells.

